# Engineering Xylose Fermentation in an Industrial Yeast: Continuous Cultivation as a Tool for Selecting Improved Strains

**DOI:** 10.1101/2022.12.13.520281

**Authors:** Thalita Peixoto Basso, Dielle Pierotti Procópio, Thais Helena Costa Petrin, Thamiris Guerra Giacon, Yong-Su Jin, Thiago Olitta Basso, Luiz Carlos Basso

## Abstract

Production of second-generation ethanol from lignocellulosic residues should be fueling the energy matrix in the near future. Lignocellulosic feedstock has received much attention as an alternative energy resource for biorefineries toward reducing the demand for fossil resources, contributing to a future sustainable bio-based economy. Fermentation of lignocellulosic hydrolysates poses many scientific and technological challenges as the drawback of *Saccharomyces cerevisiae’s* inability in fermenting pentose sugars (derived from hemicellulose). To overcome the inability of *S. cerevisiae* to ferment xylose and increase yeast robustness in the presence of inhibitory compound-containing media, the industrial *S. cerevisiae* strain SA-1 was engineered using CRISPR-Cas9 with the oxidoreductive xylose pathway from *Scheffersomyces stipitis* (encoded by *XYL1, XYL2*, and *XYL3*). The engineered strain was then cultivated in a xylose-limited chemostat under increasing dilution rates (for 64 days) to improve its xylose consumption kinetics under aerobic conditions. The evolved strain (DPY06) and its parental strain (SA-1 XR/XDH) were evaluated under anaerobic conditions in complex media. DPY06 consumed xylose faster, exhibiting an increase of 70% in xylose consumption rate at 72h of cultivation compared to its parental strain, indicating that laboratory evolution improved xylose uptake of SA-1 XR/XDH.

**GRAPHICAL ABSTRACT:** 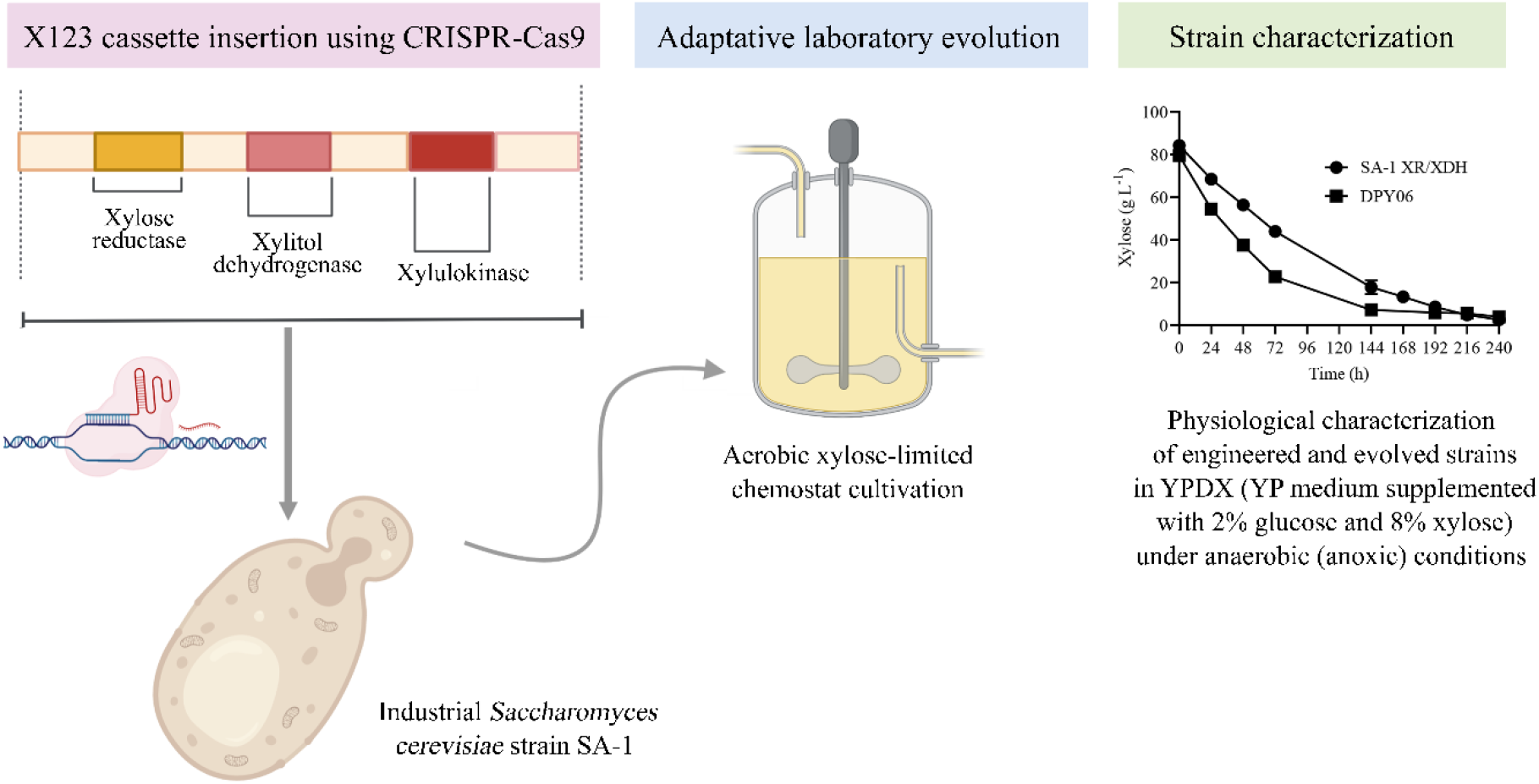

## 1. INTRODUCTION

The biomass-to-biorefinery concept is based on the replacement of first-generation feedstocks which encompasses a great spectrum of marketable products (chemicals, materials, and food) and energy (fuels, power, and/or heat) (Hassan, Williams & Jaiswal, 2019). In energy-driven biorefineries, lignocellulosic biomass is used for the production of liquid (biodiesel or bioethanol) and/or gaseous (biomethane) road transportation biofuels (Jungmeier *et al*., 2013). Lignocellulosic biomass is a potential sustainable source of various carbon sources found in several types of raw materials, ranging from urban and industrial waste, wood, and agricultural residues such as corn straw, wheat straw, rice straw, and sugarcane bagasse (Himmel *et al*., 2007; Wang *et al*., 2021). Therefore, this material represents a partially untapped rich source for biotechnology, biofuels, and biomaterials production, besides constituting a renewable alternative to petrochemicals (Guerriero *et al*., 2016). Cost-effective bioethanol production from renewable biomass requires the efficient and complete use of all sugars present in these raw lignocellulosic materials, including all pentose and hexose sugars (Li *et al*., 2015b). In particular, glucose and xylose, the major cellulose- and hemicellulose-derived sugar, respectively (Wang *et al*., 2021).

With extensive use in various biotechnological applications, including first-generation biofuels, food products, biochemicals, and the pharmaceuticals industry (Nielsen, 2019), it was natural that *S. cerevisiae* became extended toward the use in second-generation biofuel production (Li, Chen & Nielsen, 2019). However, the use of this microorganism for energy-driven biorefineries, requires that the xylose assimilation pathway is re-constructed in this microbial cell to overcome its inability to ferment xylose (Li *et al*., 2019; Procópio *et al*., 2022). For instance, this microorganism has been engineered to express either xylose reductase-xylitol dehydrogenase (XR/XDH) genes (the so-called oxidoreductase pathway), the xylose isomerase (XI) gene, or selected genes from the non-phosphorylating portion of the Weimberg pathway (Jeffries, 2006; Petrovič, 2015; Jo *et al*., 2017; Borgström *et al*., 2019; Shen *et al*., 2020; Lee *et al*., 2021).

Successfully genetic engineering allows *S. cerevisiae* strains to use xylose, however, this yeast faces uptake issues for this pentose sugar, which means that improvements in efficient xylose utilization have been hampered, in part, by xylose transport capacity (Nijland *et al*., 2014; Apel *et al*., 2016; Jiang *et al*., 2020). Despite this yeast has numerous monosaccharide transporters (*HXT1-17* and *GAL2*) only a few of those (*HXT1, HXT2, HXT4, HXT5, HXT7*, and *GAL2*) were identified to be able to transport xylose, albeit with a poor affinity that does not support growth on xylose (Sedlak & Ho, 2004). Moreover, glucose represses xylose transport by the native transporters, limiting their use in mixed sugar fermentation (Subtil & Boles, 2012; Apel *et al*., 2016; Patiño *et al*., 2019). In an attempt to overcome this impasse and allow this microorganism successfully ferment the main fraction of hemicellulose expanding bioethanol production without increasing land use (Picoli & Machado, 2021), several studies have focused on bioprospecting and characterizing heterologous xylose-transporters in *S. cerevisiae*, providing several alternatives for efficient xylose uptake (Saloheimo *et al*., 2007; Runquist *et al*., 2009; Young *et al*., 2011; Apel *et al*., 2016; Podolsky *et al*., 2021); in improving native transporters using a combination of bioinformatics and mutagenesis (Young *et al*., 2011; Farwick *et al*., 2014); and in applying adaptive laboratory evolution (ALE) to generating evolved microbial strains with desired phenotypic traits (Sonderegger & Sauer, 2003; Lee, Jellison & Alper, 2012; Parreiras *et al*., 2014; Borgström *et al*., 2019).

The experimental situation referred to as ALE are those in which the microorganisms are subjected to a variety of stressors in the environment forcing them to make evolutionary changes in their regulatory networks for optimal growth (Sauer, 2001; Mendizabal *et al*., 2014; Jeong, Lee & Kim, 2016). Chemostat culture is a method used to obtain steady-state cells and increase productivity in fermentation processes (Dragosits & Mattanovich, 2013; Jeong *et al*., 2016). Conversely, in long-term exposure to chemostat culture, the steady-state productivity is changing to the accumulation of spontaneous mutations and selection during culture under designed conditions (Wright *et al*., 2011; Jeong *et al*., 2016; Wortel *et al*., 2016). Although chemostat cultures for ALE require more complicated culture devices compared to colony transfer and serial transfer methods, it is less labor intensive once the operation begins (Dragosits & Mattanovich, 2013). Besides, chemostat culture results in the highest number of cell divisions and, therefore the highest number of diverse populations (Jeong *et al*., 2016).

Combining rational engineering followed by ALE on xylose-containing medium under aerobic (Diao *et al*., 2013) or oxygen-limited conditions (Kim *et al*., 2013b) has been successfully applied to generated *S. cerevisiae* strains with improved kinetics of xylose concerning their parental strains (Wright *et al*., 2011; Parreiras *et al*., 2014). In the present work, this strategy was chosen to expand the capabilities of an industrial *S. cerevisiae* strain for the utilization of xylose. The selected industrial strain SA-1 was engineered using a highly efficient CRISPR-Cas9 system-based approach for including the oxidoreductase pathway from *S. stipitis* (encoded by *XYL1, XYL2*, and *XYL3*) (Kim *et al*., 2013a, 2013b), yielding the SA-1 XR/XDH strain. In order to reach appropriate xylose conversion to ethanol, ALE through chemostat cultivations was used to address the low xylose uptake rate of SA-1 XR/XDH. This strain was cultivated for 64 days in an aerobic xylose-limited chemostat, resulting in the evolved strain DPY06, which showed significantly increased cell growth in xylose and glucose under anaerobic conditions.

## 2. RESULTS AND DISCUSSION

### 2.1. Construction of an industrial xylose-utilizing strain by integration of a three-gene cassette using CRISPR-Cas9 and evaluation in YPX medium

In this study, we measured the capacity of the industrial *S. cerevisiae* diploid strain, SA-1, to ferment xylose as a carbon source after expressing the oxidoreductase pathway for metabolizing xylose. Thereby, SA-1 was engineered for xylose utilization by integration through a CRISPR-Cas9-mediated system of a three-gene cassette (X123) that expresses three heterologous enzymes from *S. stipitis* necessary for xylose utilization: xylose reductase (*XYL1*), xylitol dehydrogenase (*XYL2*) and xylulokinase (*XYL3*) (Jin *et al*., 2003; Kim *et al*., 2012) into the *URA3* locus, yielding the SA-1 XR/XDH strain. Both SA-1 and SA-1 XR/XDH strains were evaluated in YPX4 at 30 °C for 72 hours, under an oxygen-limited condition in 96-well microplate equipment. Only the engineered strain, that contained the integrated xylose cassette, was able to grow in YPX under these conditions (**Figure 1**). Under this condition, the maximum growth rate (μ_max_) of the engineered strain was 0.06 h^-1^, which is higher than that presented by the xylose-consuming *S. cerevisiae* strain, BSN3, 0.044 h^-1^, when cultivated under the oxygen-limited condition in YP supplemented with 4% of xylose (Li *et al*., 2016). In their work, Li and coauthors used the xylose-consuming diploid *S. cerevisiae* strain BSIF (Li *et al*., 2015a) as the chassis cell for additional integration, including the overexpression of the genes *XKSI, TAL1*, *TKL1*, *RKL1*, and *RPE1* in the non-oxidative pentose phosphate pathway and the inactivation of *GRE3* and *PHO13*, yielding the strain BSIF (Li *et al*., 2016).

**Figure 1.**
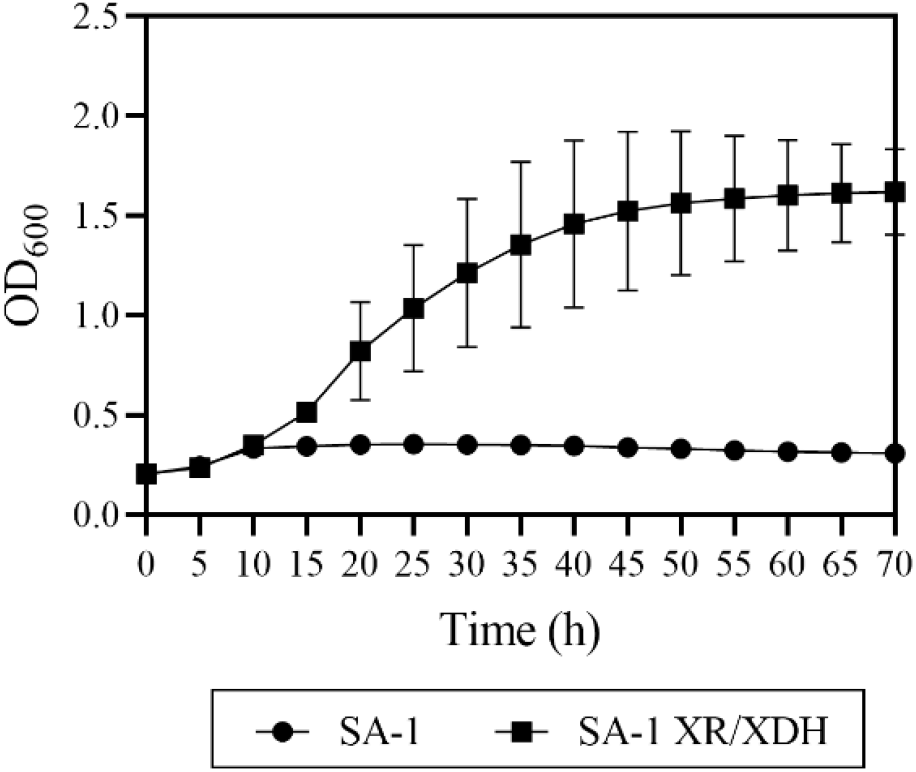
Growth kinetics for SA-1 XR/XDH yeast strain in YPX (4% of xylose) under micro-aerobic conditions at 30 °C for 72 hours in a 96-well microplate format, compared to the original strain SA-1. The graph shows the average and standard deviation for two biological replicates.

### 3.2. Physiological characterization of SA-1 XR/XDH at different xylose concentrations under anaerobic and micro-aerobic conditions

Analysis of SA-1 XR/XDH was performed in YPX medium (YP medium containing 1% xylose - YPX1, 2% xylose - YPX2, 4% xylose - YPX4, or 8% xylose - YPX8). Cells were grown under strictly anaerobic (anoxic) conditions for 624 h with an initial cell density OD_600_ of 1. The results showed that it spent more than 600 h to consume over 50% xylose when fermenting YPX1 (**Figure 2A**). In the same time frame, when fermenting 20 g L^-1^ xylose (YPX2), the SA-1 XR/XDH strain consumed 31% of total available sugar (**Figure 2B**), or 23% of 40 g L^-1^ xylose (YPX4) (**Figure 2C**), or 11% of 80 g L^-1^ xylose (YPX8) (**Figure 2D**). The increasing initial concentration of xylose did not improve xylose consumption by SA-1 XR/XDH.

**Figure 2.**
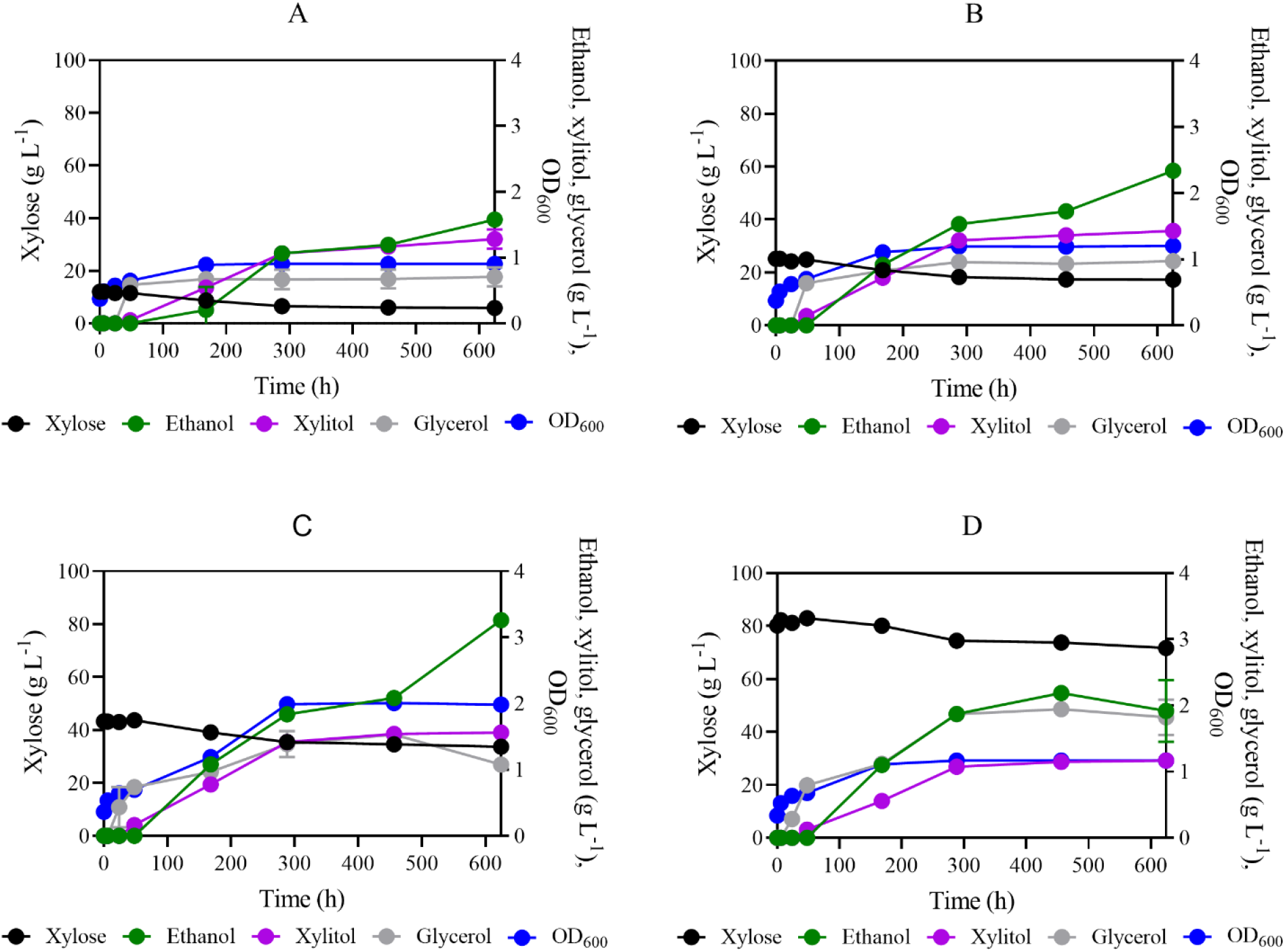
Anaerobic fermentation kinetics of SA-1 XR/XDH in YP medium containing 10 g L^-1^ xylose – YPX1 (A), 20 g L^-1^ xylose – YPX2 (B), 40 g L^-1^ xylose – YPX4 (C), or 80 g L^-1^ xylose – YPX8 (D). Fermentations were performed with initial cell density OD_600_ of 1, at 30 °C and 100 rpm.

The engineered strain achieved the highest ethanol concentration at 624 h of cultivation in the YPX4 medium (**Table 1**), which was 50%, 27%, and 34% higher for cultivation performed in YPX1, YPX2, and YPX8, respectively. Similarly, xylose metabolism rate and xylitol accumulation were higher in YPX4 cultivation. SA-1 XR/XDH (**Table 1**). Conversely, the highest maximum specific xylose consumption rate was observed for cultivations in YPX8 (0.11 ± 0.07 g xylose OD^-1^ h^-1^) within the first 24 h of cultivation. Although the rate of anaerobic xylose metabolism is relatively low, when comparing anaerobic cultivation of SA-1 XR/XDH in YPX1 with an evolved laboratory strain expressing oxidoreductase pathway from *Pichia stipitis* (Sonderegger & Sauer, 2003), the industrial xylose-fermenting strain described here achieved the same ethanol yield (0.25 g ethanol g consumed xylose^-1^) of the 460-generation population from Sonderegger and Sauer’s work.

**Table 1.** Physiological parameters of SA-1 XR/XDH in YPX1 (YP medium containing 10 g L^-1^ xylose), YPX2 (YP medium containing 20 g L^-1^ xylose), YPX4 (YP medium containing 40 g L^-1^ xylose), or YPX8 (YP medium containing 80 g L^-1^ xylose) under anaerobic conditions, at 30 °C and 100 rpm, with initial OD_600_ of 1. All parameters were calculated for up 624 h of fermentation. Parameters: Y_Ethanol_, Ethanol yield (g_ethanol_ g_consumed xylose_^-1^). Data represent the average ± standard deviation of triplicate cultivations.

| Cultivation condition | Xylose consumed (g L<sup>-1</sup>) | Xylose consumption rate (g L<sup>-1</sup> h<sup>-1</sup>) | Specific xylose consumption rate (g xylose OD<sup>-1</sup> h<sup>-1</sup>) [OD]<sup>a</sup> | Accumulated xylitol production (g L<sup>-1</sup>) | Accumulated glycerol production (g L<sup>-1</sup>) | Accumulated ethanol production (g L<sup>-1</sup>) | Y<sub>Ethanol</sub> |
| --- | --- | --- | --- | --- | --- | --- | --- |
| YPX1 | 6.21 ± 0.13 | 0.01 ± 0.00 | 0.04 ± 0.01 [0.57] | 1.21 ± 0.15 | 0.69 ± 0.15 | 1.6 ± 0.07 | 0.25 ± 0.02 |
| YPX2 | 8.23 ± 0.58 | 0.01 ± 0.00 | 0.07 ± 0.02 [0.63] | 1.43 ± 0.02 | 0.97 ± 0.03 | 2.34 ± 0.02 | 0.28 ± 0.05 |
| YPX4 | 10.39 ± 1.56 | 0.02 ± 0.00 | 0.04 ± 0.01 [0.65] | 1.56 ± 0.05 | 1.13 ± 0.09 | 3.22 ± 0.10 | 0.32 ± 0.02 |
| YPX8 | 8.59 ± 1.21 | 0.01 ± 0.00 | 0.11 ± 0.07 [0.64] | 1.16 ± 0.01 | 1.97 ± 0.27 | 2.14 ± 0.47 | 0.23 ± 0.04 |

<sup>a</sup> OD value at the maximum rate of xylose consumption.

The absence of any additional modification in SA-1 XR/XDH to address the unbalanced cofactor requirement between XR and XDH associated under anaerobic xylose fermentation resulted in the slight growth profile under strictly anaerobic (anoxic) conditions on diverse xylose concentrations (**Figure 1**). In response to the higher amount of xylose in the medium, lower xylose consumption was observed by SA-1 XR/XDH. One would think that higher xylose concentration could lead to higher xylose uptake by the yeast cell. Instead, the lower xylose assimilation by SA-1 XR/XDH for higher xylose amounts does not suggest an inefficient xylose uptake by the native transporters of the cell. These results, instead, suggest that under the anaerobic condition the highest xylose concentration (up to 40 g L^-1^) triggered catabolite repression in SA-1 yeast. In line with some previous studies of xylose-metabolizing *S. cerevisiae* which have suggested that xylose has a depressing effect on gene expression (Belinchón & Gancedo, 2003; Roca, Haack & Olsson, 2004; Salusjärvi *et al*., 2006, 2008). Anaerobic alcoholic fermentation of xylose without xylitol formation is possible when XR and XDH would have matching coenzyme specificities (Kuyper *et al*., 2004). Recent studies have demonstrated that further rational and inverse metabolic engineering strategies have improved anaerobic xylose metabolism with lesser xylitol formation of *S. cerevisiae* expressing *XYL1, XYL2*, and *XYL3* through concomitant deletion of *PHO13* and *ALD6*, as well as overexpression of *TAL1*, followed by laboratory adaptive evolution (Kim *et al*., 2013b; Jeong *et al*., 2020).

In contrast to the anaerobic (anoxic) condition, under micro-aerobic conditions, SA-1 XR/XDH could consume all xylose in 120 h of cultivation when cultivated in a YPX4 medium (**Figure 3**). Xylose consumption rate dramatically increased from 0.02 ± 0.00 g L^-1^ h^-1^ to 0.36 ± 0.00 g L^-1^ h^-1^ comparing the cultivations under anaerobic and micro-aerobic conditions, respectively, under the same cultivation medium condition (YPX4). Within the first 72 h of cultivation, SA-1 XR/XDH consumed over 90% xylose (a total of 42.49 ± 0.82 g L^-1^) producing 8.19 ± 0.34 g L^-1^ of ethanol (**Table 2**). Ethanol yield had been decreased when oxygen is available in the medium, as illustrated in **Tables 2** and **3**. Conversely, accumulated xylitol production achieved the same value for both YPX4 cultivation in anaerobic and micro-aerobic conditions. The presence of oxygen enables SA-1 XR/XDH to consume xylose faster, however, it was not enough to reduce xylitol production. Under similar conditions, SA-1 XR/XDH presented an increased xylose consumption profile than the YSX3 *S. cerevisiae* strain (Jin & Jeffries, 2004). Their strain, which contains *XYL1, XYL2*, and *XYL3* in the chromosome, was unable to consume completely 40 g L^-1^ xylose in 100 h of cultivation.

**Figure 3.**
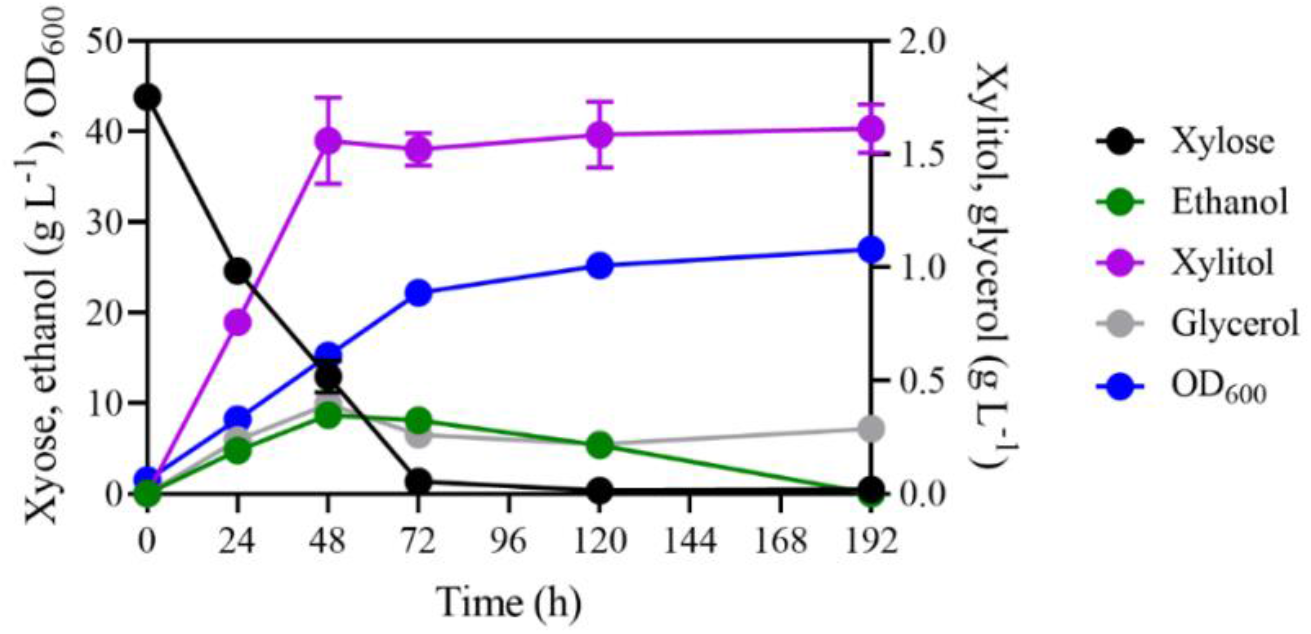
Micro-aerobic fermentation kinetics of SA-1 XR/XDH in YP medium containing 40 g L^-1^ xylose – YPX4, with initial cell density OD_600_ of 1, at 30 °C and 100 rpm.

**Table 2.** Physiological parameters of SA-1 XR/XDH in YPX4 (YP medium containing 40 g L^-1^ xylose) under micro-aerobic conditions, at 30 °C and 100 rpm, with an initial OD_600_ of 1. All parameters were calculated for up 72 h of fermentation. Parameters: Y_Ethanol_, Ethanol yield (g_ethanol_ g_consumed xylose_^-1^). Data represent the average ± standard deviation of triplicate cultivations.

| Cultivation condition | Xylose consumed (g L<sup>-1</sup>) | Xylose consumption rate (g L<sup>-1</sup> h<sup>-1</sup>) | Accumulated xylitol production (g L<sup>-1</sup>) | Accumulated glycerol production (g L<sup>-1</sup>) | Accumulated ethanol production (g L<sup>-1</sup>) | Y<sub>Ethanol</sub> |
| --- | --- | --- | --- | --- | --- | --- |
| YPX4 | 42.49 ± 0.82 | 0.36 ± 0.00 | 1.54 ± 0.01 | 0.26 ± 0.00 | 8.19 ± 0.34 | 0.24 ± 0.03 |

**Table 3.** Physiological parameters of SA-1 XR/XDH and DPY06 in YP media containing 20 g L^-1^ glucose and 80 g L^-1^ xylose under anaerobic conditions. An initial OD_600_ was adjusted to 0.3. All parameters were calculated for up 72 h of fermentation. Parameters: r_xylose_, xylose consumption rate (g L^-1^ h^-1^); r_xylose_*, specific xylose consumption rate (g L^-1^ OD^-1^ h^-1^); PEthanol, volumetric ethanol productivity (g L^-1^ h^-1^); Y_Ethanol_, ethanol yield (g g_total sugar_^-1^). Data represent the average ± standard deviation of triplicate cultivations.

|  | At 24 h |  |  | At 72 h |  |  |  |
| --- | --- | --- | --- | --- | --- | --- | --- |
| | $r_{xylose}$ | $r_{xylose}^*$ | $P_{Ethanol}$ | $r_{xylose}$ | $r_{xylose}^*$ | $P_{Ethanol}$ | $Y_{Ethanol}$ |
| **SA-1-XR/XDH** | 0.67 ± 0.07 | 0.14 ± 0.02 | 0.45 ± 0.02 | 0.56 ± 0.00 | 0.10 ± 0.00 | 0.24 ± 0.01 | 0.28 ± 0.00 |
| **DPY06** | 1.04 ± 0.01 | 0.50 ± 0.01 | 0.62 ± 0.01 | 0.79 ± 0.01 | 0.26 ± 0.00 | 0.33 ± 0.00 | 0.27 ± 0.00 |

### 2.2. Adaptive laboratory evolution of the industrial strain SA-1 XR/XDH in xylose-limited continuous cultures

In order to improve the yield and/or productivity of xylose fermentation, continuous aerobically cultivation was performed in a bioreactor. Long-term adaptation experiments are an interesting approach to selecting strains with the desired phenotype (Wortel *et al*., 2016). The ALE procedure represents one of the most practical solutions for metabolic engineering to induce spontaneous mutations, which is analogous to processes that occur in nature during the evolution of new species, favorable to heterologous metabolism, (Shin *et al*., 2021). This approach has been shown as a powerful approach for improving xylose utilization (Li *et al*., 2019). Thereby, rationally designed genetic modifications combined with ALE promote generating strains able to assimilate more efficiently xylose (Mavrommati *et al*., 2022).

After transformant evaluation, the resulting strain SA-1 XR/XDH was grown under the stress of xylose as the sole carbon source in the VM-defined medium (Verduyn *et al*., 1992). The cultivation was characterized by a batch and a continuous phase. After the batch phase, denoted by the exhaustion of xylose in the bioreactor, feeding started at a fixed dilution rate. The initial flow rate was based on the maximum growth rate of the batch phase, and the flow was kept within an interval of 8-10 mL h^-1^. Cells were cultivated at a fixed dilution rate until a decrease in xylose concentration. Therefore, as the residual xylose concentration decreased, the flow rate was augmented. Xylose consumption profile (**Figure 4.A**), glycerol, and xylitol production profiles (**Figure 4.B**) were observed during the continuous phase, but ethanol production was not.

**Figure 4.**
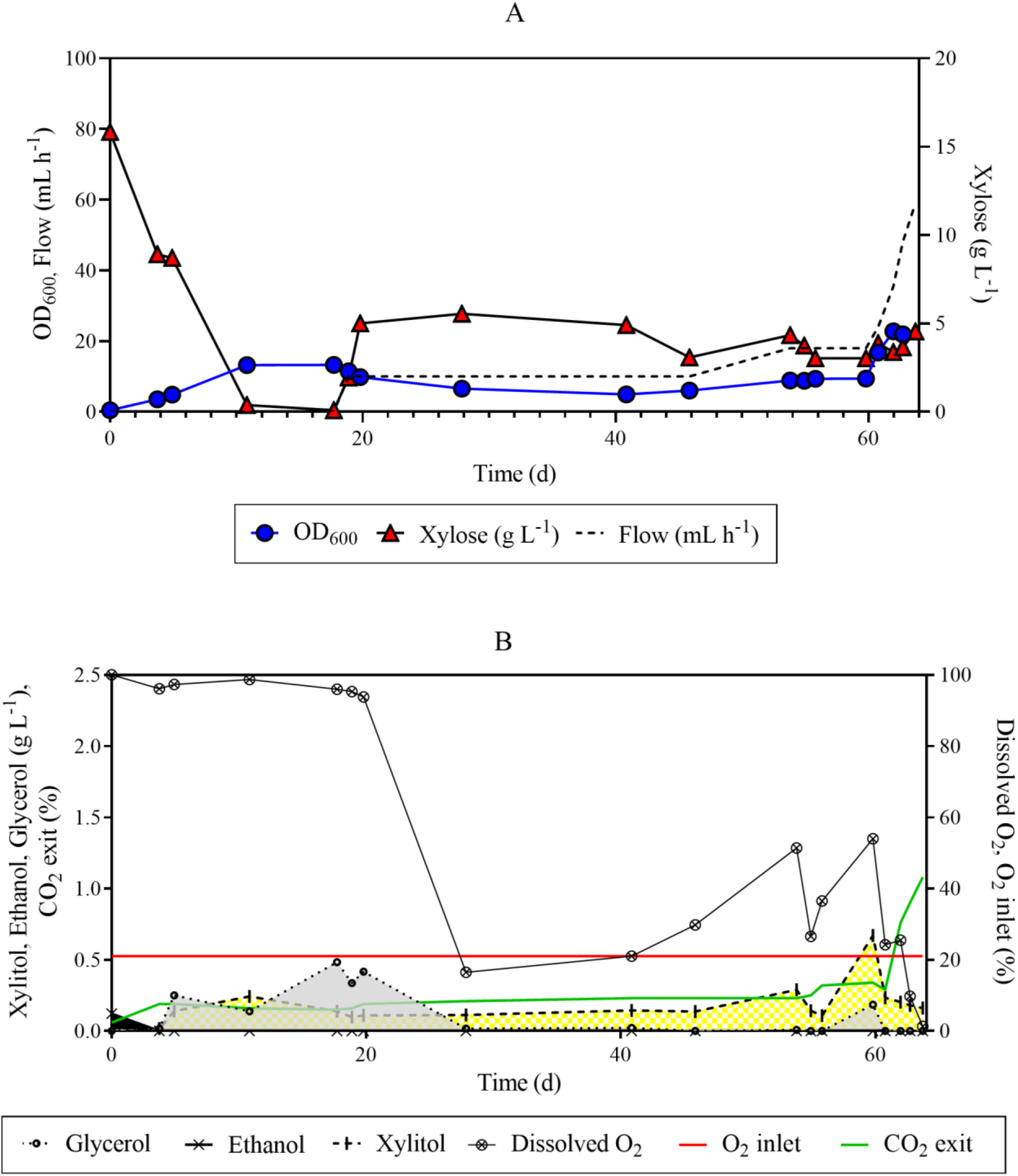
Long-term continuous cultivation of *S. cerevisiae* SA-1 XR/XDH under an anaerobic condition with xylose as the carbon source. A. OD_600_ values, residual xylose concentration, and pump flow rate. B. Glycerol, grey area; Xylitol, yellow area; Ethanol, black area. The batch phase lasted for 17 days.

After adaptive incubation on xylose, the cell density (measured by OD_600_) dropped from 13.25 at 18 h to 22.8 at 63 h of cultivation. Simultaneously, the flow of fresh medium increased from 10 to 48 mL h^-1^, corresponding to the dilution rate of 0.6 h^-1^. The increase in the flow rate did not promote the wash-out of the cells from the bioreactor, suggesting the progress of adaptive evolution, in which less adapted cells were eliminated from these cropping, remaining evolved strains with the ability to uptake xylose faster. Among the tested isolated, the colony that grew fastest on xylose was named DPY06 (data not shown).

### 2.3. Physiological characterization of SA-1 XR/XDH and the evolved strain under anaerobic conditions in xylose/glucose mixtures

To check the improvements derived from the laboratory evolution approach, the evolved strain, DPY06, and its parental strain, SA-1 XR/XDH, were cultivated under strict anaerobic (anoxic) conditions in a YP medium containing 20 g L^-1^ glucose and 80 g L^-1^ xylose, at 30 °C and 100 rpm in serum bottles sealed with butyl rubber stoppers with an OD_600_ of 0.3. DPY06 exhibited a different improved xylose consumption profile under anaerobic conditions in comparison with the parental strain (**Figure 5**). The evolved strain consumed 71% of the initial xylose in the medium, while SA-1 XR/XDH consumed only 48% in 72 h of cultivation (**Figure 5**). For glucose metabolism, no difference was observed between the two strains. However, despite the greater consumption of xylose by the DPY06 strain, the evolved strain achieved lesser cell density than the control strain, OD_600_ of 1.22 ± 0.04 versus 5.5 ± 0.16 at 192 h of cultivation. Ethanol production was detected in both cultivations, however, DPY06 did show a slight improvement in the volumetric ethanol productivity (**Table 3**). The ethanol production profile was higher for DPY06 than SA-1 XR/XDH in 144 h of cultivation, 27.1 ± 0.34 against 24.4 ±0.76, respectively (**Figure 5**). At the same time, the evolved strain produced 39% more glycerol and 9% more xylitol as a by-product than the control cultivation. Moreover, despite DPY06 showing higher xylitol accumulation until 168 h of cultivation, the evolved strain exhibited lower xylitol accumulation within 192 and 240 h of fermentation than SA-1 XR/XDH.

**Figure 5.**
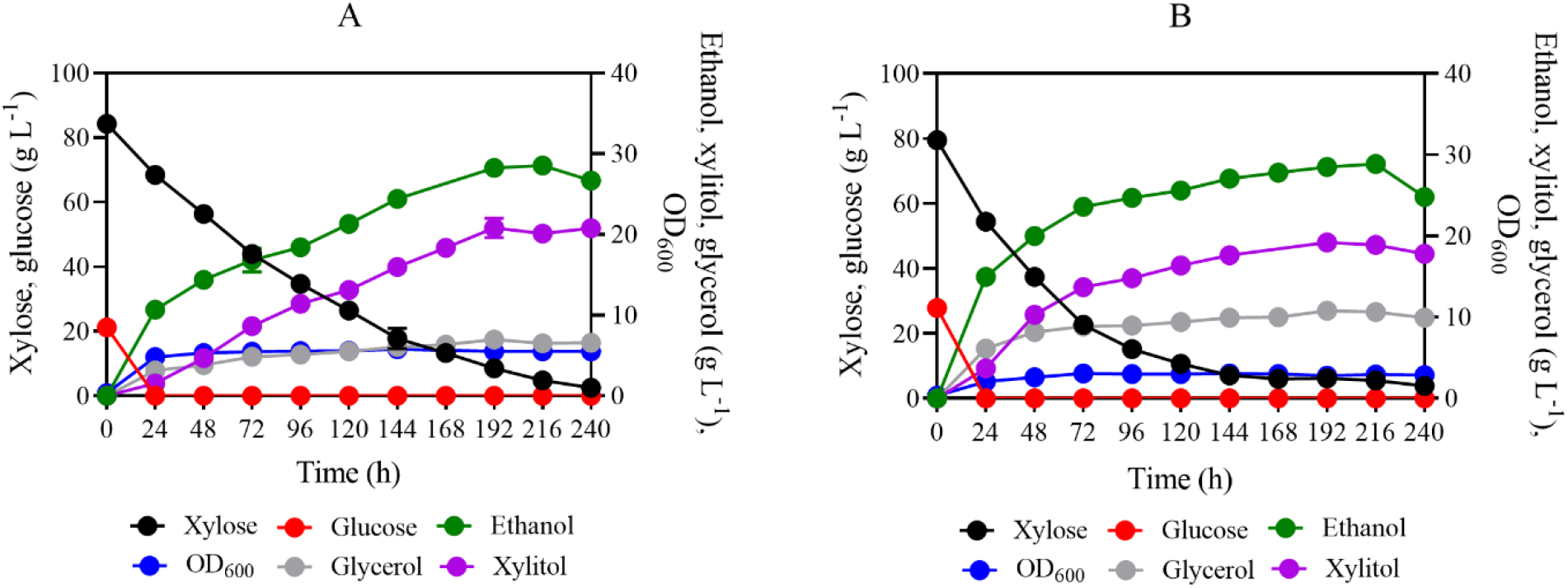
Anaerobic fermentation kinetics of the parental strain SA-1 XR/XDH (A) and evolved strain DPY06 (B) in 20 g L^-1^ glucose and 80 g L^-1^ xylose in YP media. An initial OD_600_ was adjusted to 0.3. The figure illustrates the means of triplicate experiments of each strain SA-1 XR/XDH expressing two copies of *XYL1, XYL2*, and *XYL3*; DPY06 an evolved SA-1 XR/XDH.

It is interesting to note that DPY06 used the available carbon mainly to produce ethanol, xylitol, and glycerol; and minor amounts were diverted to yeast biomass in comparison to SA-1 XR/XDH, which achieved roughly the double OD_600_ reading after 24h until the end of the cultivation. Furthermore, practically all biomass was produced during glucose depletion (first 24 h of cultivation), after that, it was no significant increments in this parameter for both strains. Xylose consumption and specific xylose consumption rates were determined and DPY06 presented high values for these parameters. **Table 3** presents these values at 24 and 72 h. Therefore, under the conditions investigated, the isolated mutant has advantages in xylose fermentation.

Similar continuous cultivation under aerobic conditions was performed by Wahlbom et al. (2003a) with recombinant xylose-utilizing *S. cerevisiae* strain TMB3399. They also performed continuous cultivation under oxygen-limited and anaerobic conditions to obtain mutants that display an improved ability to ferment xylose. However, even the best of these mutants, the strain TMB3400, showed only about one-third of the aerobic maximum growth rate obtained with *P. stipitis* CBS 6054 on xylose. Compared to its parental strain, the TMB3400 strain achieved an ethanol yield of 0.25 while for TMB3399 it was 0.21 g ethanol g xylose^-1^ (Wahlbom *et al*., 2003a, 2003b).

In a more recent study, Jeong et al (2020) applied a blended approach, metabolic and evolutionary engineering, to improve xylose assimilation in a laboratory *S. cerevisiae* strain expressing XYL123, besides a deletion of *PHO13* and overexpression of *TAL1*. An adaptive evolution approach was applied, and mutants isolated from the evolved cultures showed improved xylose fermentation capabilities, with a growth rate of 0.19 g L^-1^ h^-1^ and ethanol yield of 0.32 g g^-1^ when fermenting 40 g L^-1^ of xylose in YP medium, which was higher than DPY06 strain in YPDX under microaerobic condition. Therefore, multiple mutations were probably necessary to endow SA-1 XR/XDH with the ability to grow faster on xylose.

Although many recombinant *S. cerevisiae* strains can convert xylose into ethanol, most of these strains are haploid versions, since they are easy for genetic manipulation but, not for robustness. On the other hand, industrial diploid or polyploid *S. cerevisiae* strains are harder to manipulate but are preferred for eventual industrial applications because they are typically more tolerant to these toxic compounds and easier to cultivate in large bioreactors (Dias Lopes *et al*., 2017). These toxic compounds such as furans, organic acids, phenols, and inorganic salts are formed during lignocellulosic pre-treatment. Diploid strain presents a differentiated capacity to grow under inhibitory conditions (Cola *et al*., 2020). The genetic background of the host strain significantly affects the performance of the recombinant strain (Li *et al*., 2016). Normally, industrial diploid strains are more robust and have better ethanol producers compared to laboratory haploid strains, however, industrial diploid *S. cerevisiae* strains have been less common (Diao *et al*., 2013; Li *et al*., 2016; Cola *et al*., 2020).

Furthermore, the SA-1 strain represents a promising platform yeast strain for second-generation ethanol production, since it showed to be more robust in the presence of inhibitory compound-containing media (Cola *et al*., 2020). In the present study, the laboratory evolutionary engineering approach was a critical step to improve the xylose-fermenting capabilities of the engineered SA-1 strain.

## 3. MATERIAL AND METHODS

### 3.1. Strains and media

The yeast strain used in this study was *S. cerevisiae* SA-1, a robust industrial strain isolated from fuel ethanol plants in Brazil (Basso *et al*., 2008), which was used as microbial chassis for expressing the oxidoreductase pathway for xylose consumption from *S. stipitis*, yielding strain SA-1 XR/XDH (**Table 4**). This strain presented the genes *XYL1, XYL2*, and *XYL3* (Kim *et al*., 2012) integrated into the *URA3* locus, yielding an auxotrophic strain for uracil. Then, SA-1 XR/XDH was subjected to evolutionary engineering experiments (via adaptive laboratory evolution) in continuous cultures to improve xylose utilization, yielding strain DPY06 (**Table 4**).

**Table 4.**
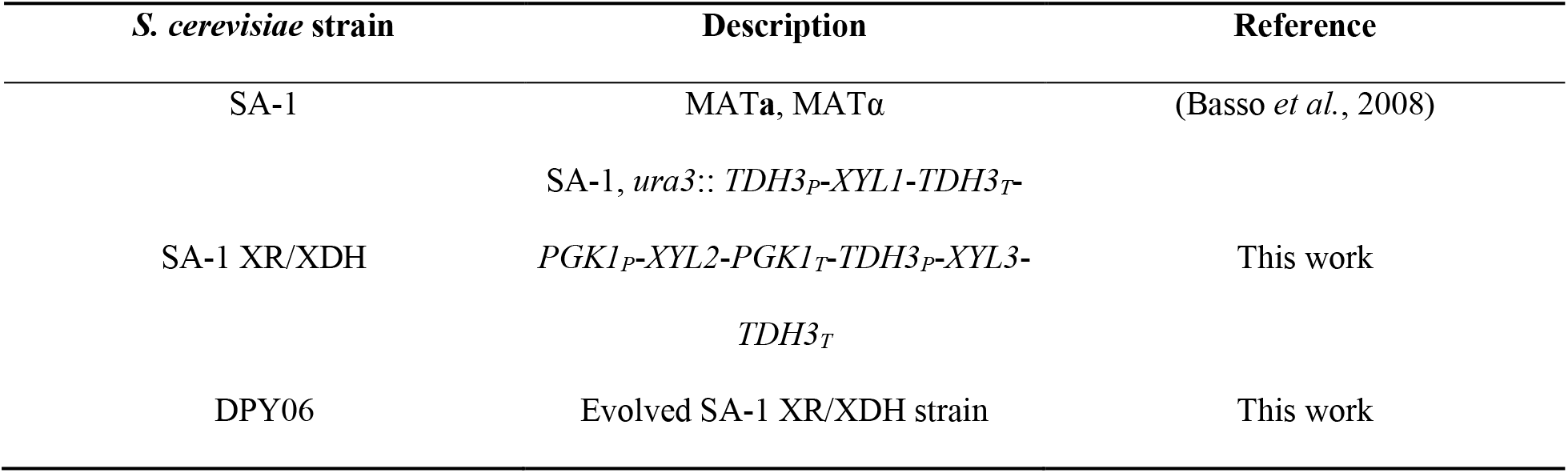
Yeast strains used in this study

Strains were cultured in media composed of yeast extract and peptone (YP, 10 g L^-1^ yeast extract, 20 g L^-1^ peptone) supplemented with glucose and or/xylose at different concentrations, both in aerobic shake-flasks and anaerobic serum-bottles batch cultivations. In adaptive laboratory evolution cultivations, strains were cultured in a defined media developed by Verduyn and co-workers (VM), as described by Verduyn *et al*. (1992). The medium was composed of the following components (in g.L^-1^): (NH_4_)_2_SO_4_, 5.0; KH_2_PO_4_, 3.0; MgSO_4_.7H_2_O, 0.5; and trace-elements consisting of (in mg.L^-1^) EDTA, 15, ZnSO_4_.7H_2_O, 4.5, MnCl_2_.2H_2_O, 0.84; CoCl_2_.6H_2_O, 0.3; CuSO_4_.5H_2_O, 0.3; Na_2_MoO_4_.2H_2_O, 0.4; CaCl_2_.2H_2_O, 4.5, FeSO_4_.7H_2_O, 3.0, H_3_BO_3_, 1.0, KI, 0.1. A solution containing vitamins was filter-sterilized and added to the medium to a final concentration of (mg L^-1^) d-biotin, 0.05; calcium pantothenate, 1.0; nicotinic acid, 1.0; myo-inositol, 25; thiamine.HCl, 1.0; pyridoxine.HCl, 1.0, and p-aminobenzoic acid, 0.20. Uracil was added to 80 mg L^-1^, as proposed by Basso et al, 2010, to avoid uracil limitation.

### 3.2. Transformation for pentose fermentation by X123 cassette insertion using CRISPR-Cas9

The three-gene cassette (X123) was integrated into the *URA3* locus using a one-step, markerless protocol mediated by CRISPR-Cas9 (Ryan *et al*., 2014). The X123 DNA (donor DNA was generated by PCR using a plasmid template pX123 was co-transformed into SA-1 along with a Cas9 plasmid (pCas9) that expresses a sgRNA with a specific 20 nucleotides (nt) sequence targeting the *URA3* locus (ACGTTACAGAAAAGCAGGCT). The 20 nt guide directs Cas9 to generate a double-stranded DNA break at the targeted locus (*URA3* in this case), which greatly increases the frequency of homologous recombination (Gietz & Woods, 2002).

### 3.3. Phenotype evaluation in YPX medium

Strain without (SA-1) and with the X123 cassette (SA-1 XR/XDH) was evaluated in YPX4 medium (YP medium containing 4% xylose) using a 96-well microplate growth assay, incubated at 30 °C with constant agitation (orbital shaking, frequency of 9.2 Hz; with an amplitude of 4.4 mm) under oxygen-limited condition; OD_600_ nm was measured every 15 minutes using Tecan Sunrise over 72 hours. The 96-well microplate was set up with 130 μl of medium and 20 μl of the fresh cell suspension to have an initial OD_600_ nm of 0.3 and 150 μl final volume.

### 3.4. Adaptive laboratory evolution in aerobic xylose-limited chemostat cultivation

After the pre-cultivation step of the SA-1 XR/XDH strain at 30 °C and 200 rpm in 500 mL shake flasks containing 100 mL of the VM-defined medium with 20 g L^-1^ xylose and 20 mg L^-1^ uracil, yeast cells were inoculated in a 2.0 L water-jacketed model Labors 5 (Infors AG, Switzerland) bioreactor containing 0.8 L of VM defined medium, with 60 g L^-1^ xylose and 80 mg L^-1^ uracil. Bioreactor batch cultivation (BBC) was conducted until the exhaustion of xylose (which was monitored by a sharp drop in the CO_2_ concentration in the off-gas analysis). After BBC, the mode of cultivation was switched to continuous through the constant addition of fresh VM medium supplemented with 20 g L^-1^ xylose and 80 mg L^-1^ uracil by a peristaltic pump. The working volume was kept constant by a mechanical drain and a peristaltic pump in line with an aseptic waste vessel. The feeding rate was adjusted during the cultivation to increase the selective pressure on cells (please, see details in the “Results” section). Compressed air was used to flush the culture vessel (0.5 L min^-1^). The dissolved oxygen concentration was monitored constantly by a dissolved oxygen electrode (Mettler-Toledo, Columbus, OH, USA). The agitation frequency was set to 800 rpm, the temperature was controlled at 30 °C, and pH was controlled at 5.0 via a controlled 2 M KOH solution. Continuous cultivation was performed for 64 days. To isolate potential evolved strains, samples were plated from the continuous cultivation medium in a YP medium containing 20 g L^-1^ xylose (YPX2) and evaluated under aerobic (oxic) and anaerobic (anoxic) conditions (200 rpm) in shake-flask cultivations, as depicted below.

### 3.5. Shake-flask cultivations for strain characterization

YP medium was used for shake-flasks batch cultivations. After pre-inoculum, yeast cells were harvested by centrifugation at 3,134 × g, at 4°C for 5 min, and washed three times with distilled water. Aerobic and micro-aerobic batch fermentation experiments were performed by inoculating the pelleted yeast cells in a 100 mL Erlenmeyer flask with 30 mL containing fermentative medium, composed of YPX (YP medium containing xylose), YPD (YP medium containing glucose), or YPDX (YP medium containing glucose and xylose). All conditions were performed at 30 °C. For aerobic conditions, agitation was set at 200 rpm, while under anaerobic and micro-aerobic conditions, agitation was at 100 rpm. For anaerobic batch fermentation experiments, pre-cultures were inoculated in a serum bottle containing 30 mL of fermentative medium, which could be YPX or YPDX. A serum bottle sealed with butyl rubber stoppers was used to ensure strict anaerobic (anoxic) conditions. The serum bottles with fermentation media were then flushed with nitrogen gas, previously passed through a heated, reduced copper column to remove the trace of oxygen. All cultivations were performed in biological triplicates.

### 3.6. Analytical methods and calculation

Samples were taken at appropriate intervals to measure cell growth and metabolites. Cell growth was monitored by optical density (OD) at 600 nm using a UV-visible Spectrophotometer (BIomate 5) (Basso *et al*., 2010). Samples were centrifuged at 15000 rpm for 10 min and supernatants were diluted appropriately and then used for the determination of glucose, xylose, xylitol, glycerol, succinate, acetic acid, and ethanol by high-performance liquid chromatography (Agilent Technologies 1200 Series) equipped with a refractive index detector. The Rezex ROA-Organic Acid H+ (8%) column (Phenomenex Inc., Torrance, CA) was used and the columns were eluted with 5 mM H_2_SO_4_ at 50 °C, and the flow rate was set at 0.6 mL/min.

The maximum growth rate was obtained by plotting the natural logarithm of OD_600_ values against time and then calculating the slope of the straight line corresponding to the exponential growth phase. Yield coefficients were calculated as the absolute value of the slope of a straight line: the ethanol yield (Y_ethanol_) from a plot of product concentration data against substrate concentration data (Della-Bianca & Gombert, 2013)

## 5. CONCLUSIONS

Laboratory adaptive evolution provided an improvement in SA-1 XR/XDH strain, mainly concerning the xylose uptake. This dataset demonstrates the feasibility of this strategy. The evolved strain, DPY06, was able to consume xylose faster, presenting a specific xylose consuming rate 72% higher than the control cultivation, and showed an improvement in the volumetric ethanol productivity, 28% comparing DPY06 and SA-1 XR/XDH at 24 h of cultivation in YP media supplemented with glucose and xylose. The evolutive study was performed in long-term continuous cultivation in synthetic media under controlled conditions. Although we have verified that the evolved strain performs well at elevated sugar concentrations (100 g L^-1^ sugars) under strictly anaerobic conditions, many other parameters relevant for industrial second-generation ethanol production remain to be systematically investigated, as full incorporation of lignocellulosic hydrolysates into the fermentative analysis of studied strain.

## 6. ACKNOWLEDGMENT

We thank MSc. Gabriel C. de Gois e Cunha for helping with analytical methods, and Dr. Bruno L. V. da Costa for assisting with bioreactor operation. We are also grateful to Dr. Jeffrey M. Skerker and Prof. Adam P. Arkin for guidance on engineering yeast strain SA-1 XR/XDH. This research was funded by Fundação de Apoio à Pesquisa do Estado de São Paulo (FAPESP) grants #2015/50612-8, #2018/17172-2, #2018/01759-4, and #2019/18075-3.

## 7. CREDIT AUTHORSHIP CONTRIBUTION STATEMENT

**Thalita Peixoto Basso:** Methodology, Investigation, Data curation. **Dielle Pierotti Procópio:** Methodology, Investigation, Formal analysis, Supervision, Data curation, Validation, Writing – original draft, preparation. **Thais Helena Costa Petrin:** Methodology. **Thamiris Guerra Giacon^-^:** Methodology. **Yong-Su Jin:** Supervision, Formal analysis. **Luiz Carlos Basso:** Formal analysis. **Thiago Olitta Basso:** Supervision, Project administration, Formal analysis, Resources, Project administration, Writing – review & editing.

